# Surviving in high stress environments: Physiological and molecular responses of lobe coral indicate nearshore adaptations to anthropogenic stressors

**DOI:** 10.1101/786673

**Authors:** Kaho H Tisthammer, Emma Timmins-Schiffman, Francois O Seneca, Brook L Nunn, Robert H Richmond

**Author notes:** Corresponding Author /.

## Abstract

Corals in nearshore marine environments are increasingly exposed to reduced water quality, which is the primary local threat to Hawaiian coral reefs. It is unclear if corals surviving in such conditions have adapted to withstand sedimentation, pollutants, and other environmental stressors. Lobe coral populations from Maunalua Bay, Hawaii showed clear genetic differentiation between the ’polluted, high-stress’ nearshore site and the ’lower-stress’ offshore site. To understand the driving force of the observed genetic partitioning, reciprocal transplant and common-garden experiments were conducted to assess phenotypic differences between these two populations. Physiological responses were significantly different between the populations, revealing more stress-resilient traits in the nearshore corals. Changes in protein profiles between the two populations highlighted the inherent differences in the cellular metabolic processes and activities; nearshore corals did not significantly alter their proteome between the sites, while offshore corals responded to nearshore transplantation with increased abundances of proteins associated with detoxification, antioxidant defense, and regulation of cellular metabolic processes. The response differences across multiple phenotypes between the populations suggest local adaptation of nearshore corals to reduced water quality. Our results provide insight into coral’s adaptive potential and its underlying processes, and reveal potential protein biomarkers that could be used to predict resiliency.

## INTRODUCTION

Coral reefs are among the most productive ecosystems on the planet, providing important benefits to diverse species that inhabit them and sustaining the lives of over 500 million people through their economic, cultural, physical, biological, and recreational services^1^. Despite their importance, coral reefs worldwide are highly threatened by local and global stressors resulting from human activities. Rates of current environmental change are orders of magnitude faster than those of ice-age transitions^2^, so the fate of coral reefs will ultimately depend on whether corals and their ecosystems can adapt or acclimatize with at a fast enough rate to mitigate rapid environmental changes. Thus far, we already observe that coral cover around the world has declined over 50% in the past 100 years^3^. Although global climate change is viewed as the dominant threat to coral reefs, localized anthropogenic stressors, such as overfishing, pollution, and coastal development, play significant roles in the decline of coral reefs^4^. Because coral reefs experiencing multiple stressors exhibit lower ecosystem resilience (e.g.^5,6^), understanding the effects of local stressors and coral’s adaptability to such stressors is vital to developing and implementing effective management interventions as global-level stressors continue to increase^4^.

Reduced water quality due to human actions is an increasing threat to nearshore marine habitats and is one of the major local threats to coral reefs^4^, especially in Hawaii. Maunalua Bay, Oahu hosts a diverse ecosystem dominated by coral reefs where environmental changes to physical and chemical water properties are well-documented and characterized^7–10^. In the last century, the health of these coral reefs has drastically deteriorated due to large-scale urbanization: Coral cover over most reef-slopes is < 5%, down from 50% in the 1950’s^8^. Corals in Maunalua Bay nearshore areas, especially in the inner bay, are under chronic stress from sedimentation and pollutant-laden terrestrial runoff^11^, yet despite prolonged exposure to these stressors, some individuals continue to survive, suggesting they may have acclimatized or adapted to withstand such stressors. Population genetic analyses also revealed clear genetic differentiation between the nearshore (N) and offshore (O) populations in Maunalua Bay (Fig. 1)^12^. Because the distance between the two sites is small (< 2 km) with no apparent barriers^10^, the results suggest local selection as the driving force of the observed genetic partitioning^12^.

**Figure 1.**
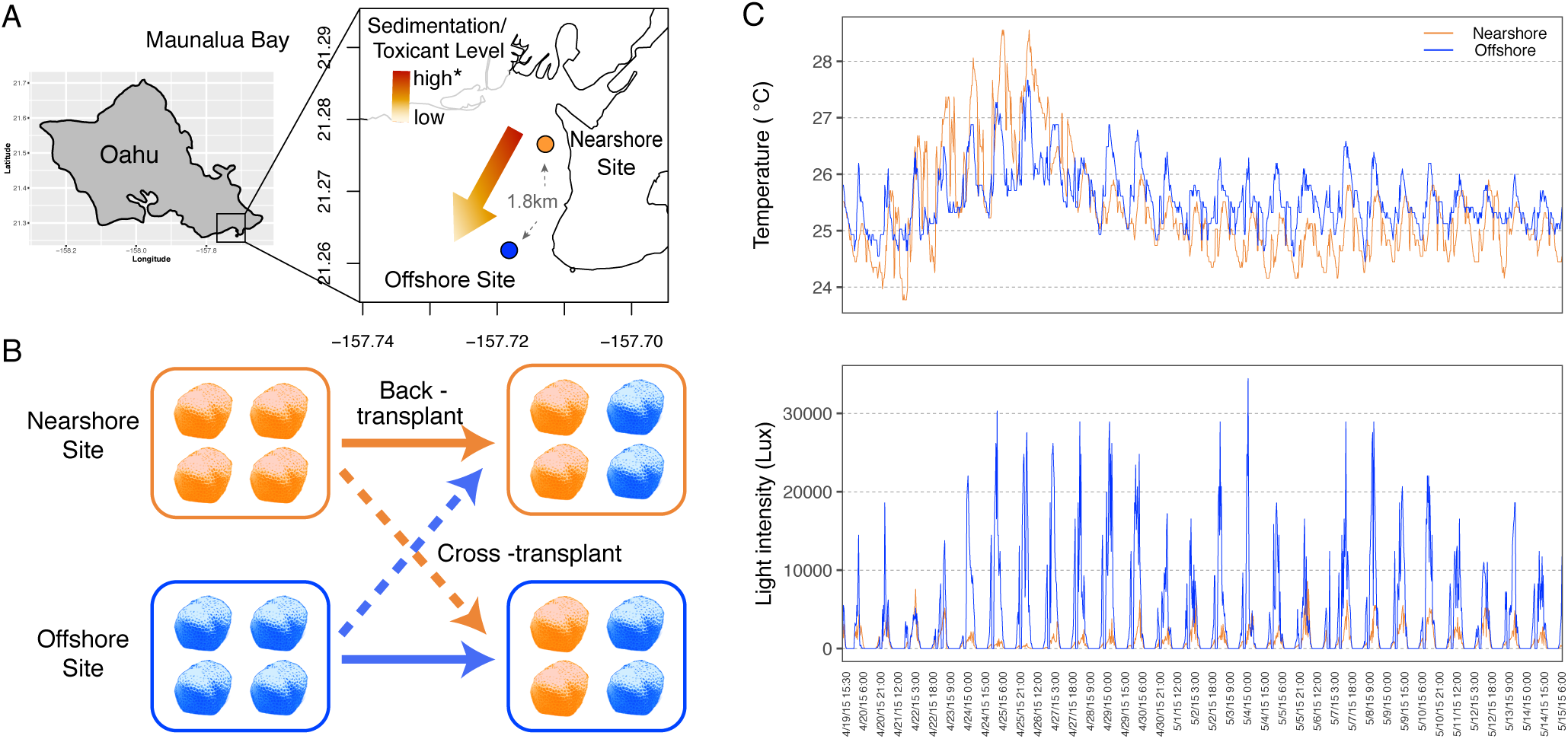
Experimental location and design. (A) a map of study locations, (B) a diagram of the reciprocal transplant experimental design, and (C) temperatures and light intensity profiles during the reciprocal transplant experiment of N- and O-sites. N-site is located in the inner bay where it receives direct run-offs, while O-site is located at the end of the bay, exposed to offshore currents with its environmental conditions closer to those of the offshore, despite its proximity to the shoreline (for detailed water quality profiles, see [8,10]).

Based on the genetic results, we tested whether N- and O-populations of *P. lobata* in Maunalua Bay was due to local adaptation using reciprocal transplant and common garden experiments. *Porites lobata* in Maunalua Bay offers a unique opportunity to study the coral’s adaptability because of accessibility and extensive environmental research history. In addition, since *P. lobata* primarily harbors a specific clade of symbiotic *Cladocopium* (C15), which is vertically transmitted with high fidelity^13^, the assessment of the coral-host’s adaptive abilities, as opposed to those of its endosymbiotic zooxanthellae, can be easily monitored. To date, no shuffling of zooxanthellae in *P. lobata* has been reported. The goal of this study was to investigate how molecular and physiological responses differed between the N-*vs.* O-populations under the divergent water-quality conditions, and to obtain any insight into the metabolic processes involved in stress tolerance of corals to reduced water quality. Analyses of tissue layer thickness, tissue lipid content, and proteomic profiles of *P. lobata* were completed following a 30-day reciprocal transplant experiment and short-term growth rates were compared using a common-garden experiment.

## RESULTS

### Physiological response differences between the populations

Small fragments from five source colonies from the two experimental sites (N- and O-sites) were used to conduct a reciprocal transplant experiment in the Maunalua Bay, Hawaii (Fig. 1). The results revealed clear physiological response differences between the two populations. The transplantation resulted in a significant reduction in the average tissue layer thickness (TLT) in only one treatment; O-corals transplanted to N-site (O→N) (Tukey-HSD, *P*-adj < 0.001, Fig. 2A, SI.1A). Contrary to the expectation of coral lipid content being reduced by environmental stress, the total tissue lipid content in O-corals showed a marginally significant increase when transplanted to N-site (Tukey-HSD, *P* = 0.059, SI.1A), while N-corals showed very little change in their lipid content between the sites (Fig. 2B). Two-way ANOVA showed no population or transplant-site effects, but a significant interaction between the two factors (*P* = 0.034). Comparing lipid contents pre- and post-experiment revealed that lipid contents of O-corals increased significantly at both sites, while those of N-corals did not (Fig. 2D). The short-term growth rates assessed using a common-garden experiment revealed a significantly higher average growth rate in N-corals than O-corals. N-corals grew on average 5.57% of their initial weight over 11 weeks, while O-corals grew 2.57% (Fig. 2C, ANOVA, *F* = 13.09, *P* = 0.0068).

**Figure 2.**
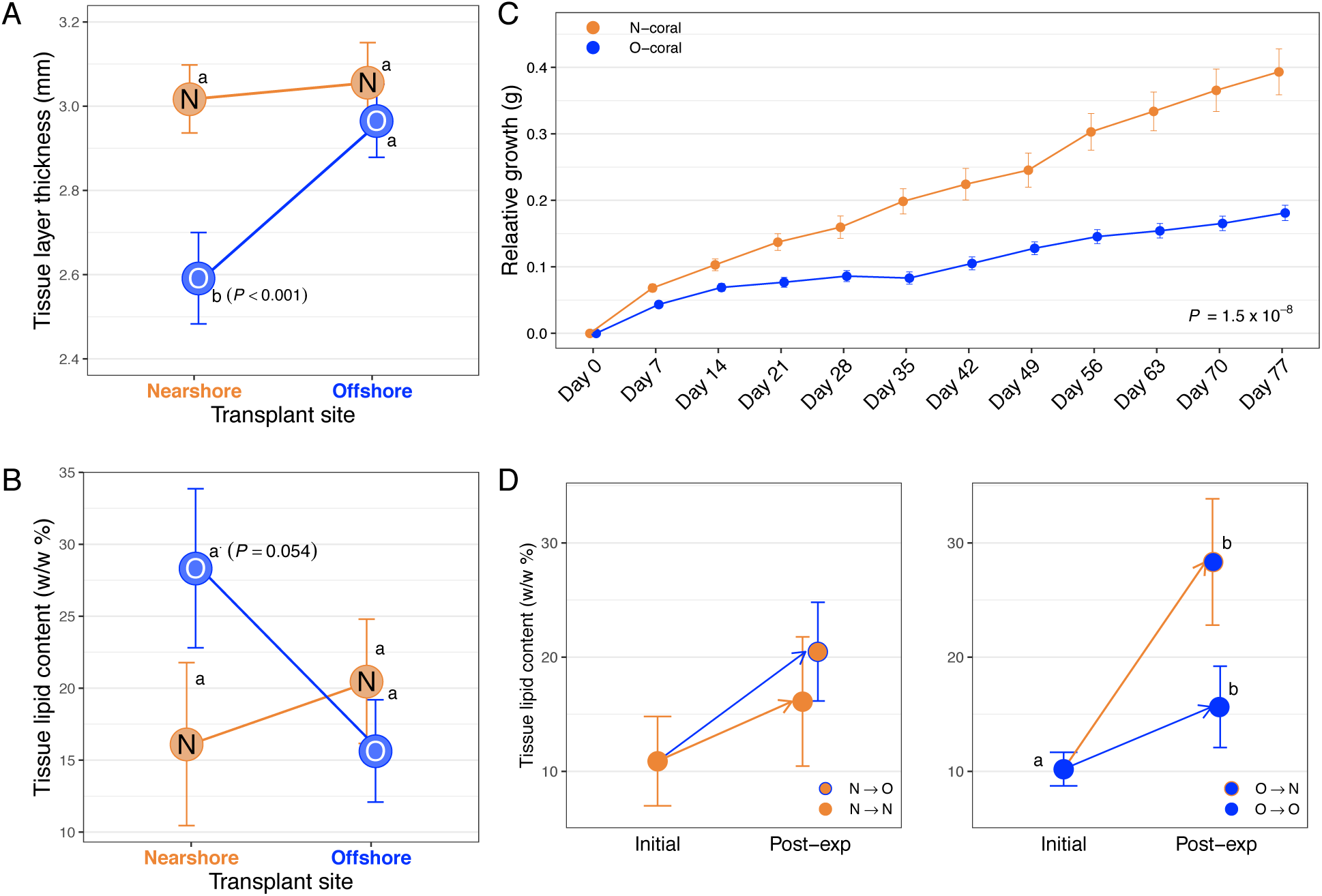
Physiological results of the 30-day reciprocal transplant experiment of *P. lobata* from N- and O-sites. (A) tissue layer thickness, (B) tissue lipid content, and (C) short-term growth rate (starting weight adjusted to 6 g), (D) tissue lipid content before and after the experiment. Different letters indicate a statistically significant difference. Error bars denote standard error.

### Transplantation effects on proteomic profiles

A label-free shotgun proteomics approach was performed to study the dynamic protein-level response to transplantation. Combining all proteomic results, a total of 3,635 proteins were identified with high confidence (FDR < 0.01) when correlated to a customized *P. lobata* protein database predicted from the transcriptome ^14^. An average of 2,777 proteins were identified across all biological replicates in each treatment. A total of 1,977 proteins were shared across all four treatments, which likely represents the basic homeostatic functions of corals. Proteins unique to a treatment ranged from 105 to 233 (Table S1, Fig. S1A).

To evaluate the differences revealed in the protein-level responses among the treatments, normalized spectral abundance factor (NSAF) values for each protein were visualized using non-metric multidimensional scaling (NMDS). NMDS analysis revealed a clear separation of the protein abundance profiles between the two populations regardless of the transplant sites (ANOSIM, *R* = 0. 7074, *P* = 0.004, Fig. S2). Transplantation appears to have had a greater effect on the final protein profile of O-corals, while protein abundances of N-corals were relatively similar between the transplant sites.

### Response differences in transplant to Nearshore site (N→N vs. O→N)

A total of 3,290 distinct coral proteins were identified in at N-site: 2,365 (72%) were shared between N→N and O→N corals, with 402 proteins unique to N→N corals, and 523 unique to O→N corals (Fig. S1B). Gene Ontology (GO) enrichment analysis (CompGO) ^15^, identified 19 enriched GO terms specific to N→N corals and 42 terms specific to O→N corals from the three GO categories (Biological Process [BP], Molecular Function [MF], and Cellular Component [CC]) (SI.2A). Enriched terms in N→N corals included peptidase inhibitor activity, oxidoreduction coenzyme metabolic process, lyase activity, and regulation of protein polymerization. The enriched terms for O→N corals included detoxification, antioxidant activity, lipid oxidation, intracellular protein transport, tricarboxylic acid (TCA) cycle, and purine-containing compound metabolic process.

Quantitative analysis on the protein level identified 138 proteins significantly more abundant (referred to as the abundant-proteins) in N→N corals and 276 in O→N corals (Fig. 3A, Table S1). GO analysis identified a total of 35 terms from the abundant-proteins in O→N corals (SI.2A), while none met the statistical cutoff in N→N corals. The enriched terms in O→N corals indicated that the abundant-proteins were dominated by those associated with amino acid metabolic process, oxidation-reduction process, and vesicle membrane/coat (SI.2A). Abundant proteins identified in O→N corals further suggested more activities in metabolic and regulatory pathways, including detoxification and glutathione pathways (*i.e.* antioxidant activity) (Fig. S3). The three most abundant proteins (with annotation) in N→N corals were associated with immune function (hemicentin-2: m.9723, glycoprotein 340: m.13233, and lectin MAL homologue: m.12716), while those in O→N corals were involved with detoxification function (arsenite methyltransferase:m.16246, E3 ubiquitin-protein ligase TRIM71:m.2139, and glutathione-S-transferase:m.16994).

**Figure 3.**
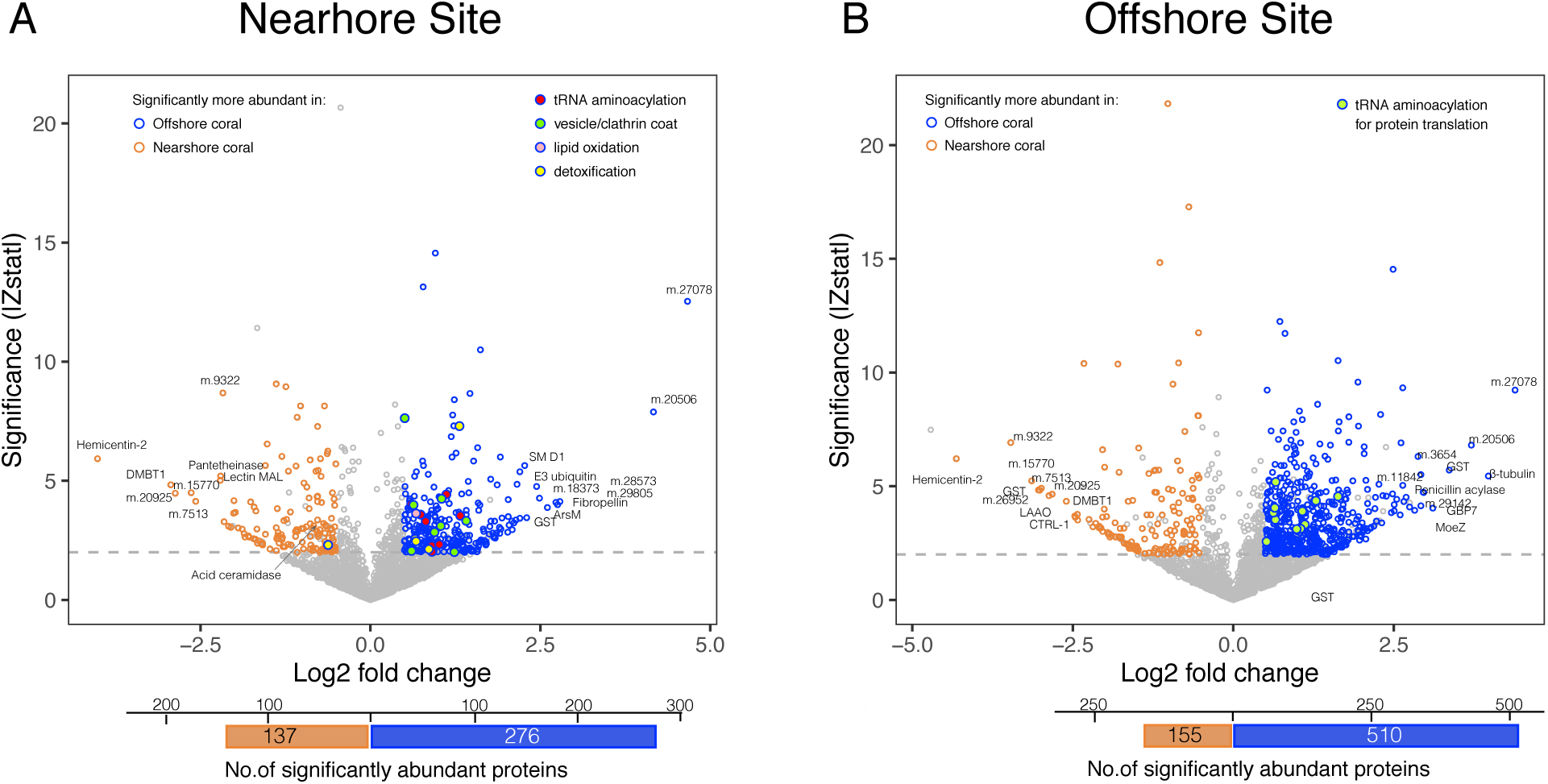
Protein abundance comparison between N- and O-corals. (A) at N-site, and (B) at O-site. Fold differences and Z-statistic values (Z-stat) were used to generate volcano plots. Colored points indicate proteins with |fold| > 0.5, |Z-stat| > 2 at FDR=0.01. Proteins associated with key GO terms were colored in different colors, and the top 10 abundant proteins in each population are annotated. The bottom bars indicate the total numbers of significantly abundant proteins for each population.

### Response difference in transplant to the Offshore Site (N→O vs. O→O)

A total of 3,236 distinct coral proteins were identified at O-site: 2,217 (68.5%) were shared between the two populations, 656 unique to N→O corals, and 363 to O→O corals (Fig. S1C). GO analysis identified 35 enriched terms specific to N→O, which involved amino acid biosynthetic process, ATP metabolic process, TCA cycles, fatty acid oxidation, and monosaccharide metabolic process. There were 15 specific GO terms in O→O corals, including nucleotide monophosphate biosynthetic process, intracellular protein transport, vesicle organization, and GTP binding (SI.2B).

Quantitative analysis on protein abundances indicated a total of 665 proteins to be significantly differentially abundant at O-site: N→O corals had 155 abundant-proteins, and O→O corals had 510 abundant-proteins (Fig. 3B). GO analysis resulted in identifying 39 enriched terms from abundant proteins in O→O corals, while only one met the cutoff in N→O corals (SI.2B). Although the number of abundant-proteins and enriched terms identified in O→O corals were relatively high, the enriched terms predominantly consisted of cellular functions related to protein translation; organonitrogen biosynthetic process and organic acid metabolic process, both leading to single child terms for BP, CC, and MF (tRNA aminoacylation for protein translation, cytosolic large ribosomal subunit, and tRNA aminoacyl ligase activity). The enriched term in N→O corals was a non-specific term of ‘extracellular region’, indicating that despite the higher number of abundant-proteins, the main functional difference between N→O and O→O corals was an enhanced protein translation activity in O→O corals.

### Response comparisons to cross transplantation

Effects of cross transplantation yielded a more diverse proteomic stress-response in O-corals as they moved nearshore than N-corals as they were moved offshore (Fig. S2). The total number of abundant-proteins between the sites was much higher for O-corals (440, O→N *vs*. O→O) than N-corals (135, N→N *vs*. N→O) (Table S1), and the number of unique GO terms identified between the sites was also higher in O-corals (69, SI.2C) than in N-corals (46, SI.2D). The number of overlapping proteins between the sites was lower in O-corals than in N-corals (70% *vs.* 79%), and log-fold changes of all identified proteins between the sites were significantly larger for O-corals than N-corals (Wilcoxon Rank-Sum test, *P* = 6.02 x10^−9^), all emphasizing the larger metabolic reshuffling needed to respond to cross transplantation in O-corals. GO enrichment analysis indicated that N-corals responded to transplantation to O-site with increased abundance in proteins involved in amino acid biosynthesis, fatty acid beta oxidation, TCA cycle, chitin catabolism, coenzyme biosynthesis and translational initiation. O-corals responded to transplantation to N-site by increasing the abundance of proteins associated with detoxification, antioxidant activity, protein complex subunit organization, and multiple metabolic processes (amino acid, fatty acid, ATP, monosaccharide, and carbohydrate derivative) (SI.2E). The shared responses between the cross-transplanted corals (N→O and O→N corals) included increased proteins involved in fatty-acid beta oxidation, TCA cycle, carbohydrate derivative catabolic process, pyridoxal phosphate binding, and ’oxidoreductase activity acting on the CH-CH group of donors with flavin as acceptor’, likely representing the effects of transplantation to a non-native environment.

### Proteome patterns across the four treatments

Comparing enriched GO terms across all treatments (SI.2E) highlighted the unique state of O→N corals; O→N corals had a much higher number of uniquely enriched GO terms (n=27) compared to those in the rests (4 in O→O, 5 in N→N, and 15 in N→O corals). The most notable difference among the treatments was enrichment of detoxification and antioxidant activity exclusively in O→N corals (Fig. 4). Also, lipid oxidation was highly enriched in O→N corals with four terms associated to this category identified (Fig. 4, SI.2E).

**Figure 4.**
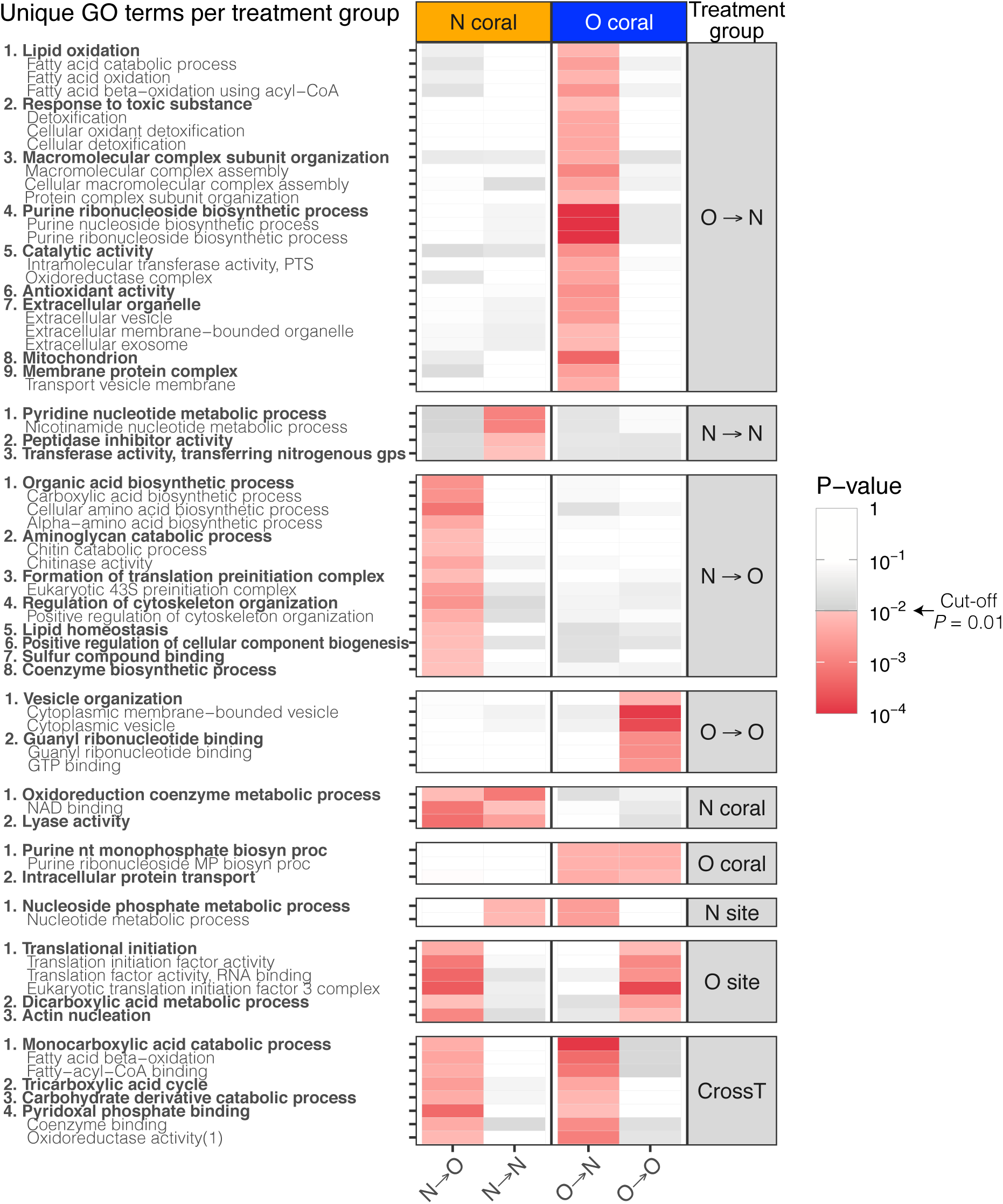
Enriched GO terms uniquely identified to specific treatment groups. Treatment groups are shown in the right column (*e.g.* N-coral = N-corals at both sites, N-site = N- and O-corals at N-site, CrossT = cross transplantation). The heat-map represents *P*-values for the associated GO terms. The GO terms are grouped by the parent-child terms with the most parent term in bold (for values, see SI-2E).

Examining the relative abundance of individual proteins associated with detoxification (’detox-proteins’) revealed the following interesting patterns. 1) Distinct sets of proteins were abundant in different treatments, rather than all detox-proteins to be elevated in one treatment, and the direction and magnitude of responses to transplantation were protein specific and varied between populations (Fig. S4A). 2) Two peroxiredoxin (Prx) proteins, Prx-1 (m.6147) and Prx-6 (m.9595), dominated the relative abundance of detox-proteins by having over an order of magnitude higher abundance values, and they were consistently more abundant in N-corals than O-corals (ave. 44%, Kruskal Test, *P* = 0.004 −0.01) (Fig. S4B, SI.1B). 3) Some proteins with the same or similar annotations had contrasting responses between the populations. For example, Prx-4 (m.17739), which belongs to the same subfamily as Prx-1, was significantly more abundant in O-corals at both sites (Fig. S4B, SI.2F;2G), while Prx-1 was more abundant in N-corals. Similarly, seven peroxidasin (PXDN) homologs were identified, of which m.17686 was significantly more abundant in O→N corals, while m.9432 was significantly more abundant in N→N corals (Fig. S4B, SI.2F), suggesting that the two populations potentially utilize different class/kind of enzymes as primary proteins in detoxification/antioxidant pathways. Of the seven PXDN homologs, two (m.1440, m.9432) were consistently higher in N-corals, two (m.10928, m.15200) were consistently higher in O-corals, and three (m.12572, m.17686, m.9657) increased abundance at N-site in both corals, but m.12572 and m.17686 being higher in O-corals, while m.9657 higher in N-corals (Fig. S3B).

To ascertain that the proteins with the same annotations are indeed different proteins, sequences of matched peptides were assessed for those that showed contrasting responses. The pairwise comparison of Prx-1 and Prx-4 showed only seven of the total 65 peptides (11 %) were identical between the two, revealing that these protein sequences are significantly different and they each have unique peptides that be detected and quantified accurately (SI.1C1). Similarly the majority of PXDN-like proteins identified had no overlapping peptides between the contrasting pairs (0 - 19%, median = 0, SI.1C2), indicating that corals possess multiple types of PXDN, and N- and O-corals respond to stressors with different sets of PXDN.

In addition to lipid oxidation being significantly enriched in O→N corals, a single term (fatty acid beta-oxidation,) was also enriched in N→O corals, which suggests that cross-transplantation had an effect on lipid oxidation processes. However, the abundances of most proteins associated with lipid oxidation were higher in O-corals than N-corals at both sites (Fig. S4A). Statistically, three proteins (medium-chain sp acyl-CoA:m.22274, very-long-chain sp. acyl-CoA:m.17984, and trifunctional enzyme subunit alpha:m.6724) showed a difference in abundance between the two populations at N-site (Fig. S4C) and one (isovaleryl-CoA dehydrogenase:m.27714) at O-site, all of which were higher in O-corals than N-corals.

## DISCUSSION

Physiological and molecular analyses of the N- and O-coral populations exposed to the same environmental conditions revealed dynamic and divergent metabolic responses. This, together with clear genetic differentiation observed between the two populations^12^, suggests that some adaptive differences exist between the two populations, although the effects of acclimatization cannot be ruled out completely. The response differences in TLT (significant reduction in O→ N corals) and growth-rate (lower in O-corals) indicate that N-corals possessed more resilient traits in response to the nearshore environmental stressors. The TLT of *Porites* corals is known to be reduced by sedimentation and other environmental stressors (e.g.^16 17^). Between our experimental sites, sedimentation/turbidity was the most variable environmental parameter. On average, the mean turbidity varies by an order of magnitude between the sites^9^, with differences being more pronounced during and after storms at N-site^8^. Much lower light intensity recorded at N-site during our experiment also suggests higher turbidity there than at O-site (Fig. 1). Therefore, the observed reduction in TLT in O→N corals was likely due to sedimentation stress, which in turn reflects higher resilience of N-corals to such a stressor. Faster growth rates observed in N-corals from the common-garden experiment also suggest potential resilience to reduced water quality, since this experiment was conducted in conditions resembling N-site. Seawater used for the experiment came from the Kewalo Channel, which receives several high-volume terrestrial discharges and boat-traffic based fuel contaminants. Additionally, the degree of daily temperature fluctuation of the experimental tank (2.97 °C daily max) mimicked N-site variability (2.83 °C) more closely than O-site variability (2.14 °C) (Fig. 1).

Increases in tissue lipid content (w/w%) observed in O→N corals were unexpected, as decline in lipids is generally reported under stressful conditions (e.g.^18^). High sedimentation, however, has been observed to alter corals’ metabolism by increasing the energy gains from heterotrophic sources^18–20^. For example, when the coral *Stylohora subseriata* was transplanted to a eutrophic, nearshore site, their tissue lipid content increased, which was hypothesized to have resulted from an increase in both heterotrophic and phototrophic feeding^21^. *Porites* species, however, show little evidence of having an ability to increase their heterotrophic feeding rate to meet their daily metabolic energy requirements: less than10% of *P. cylindrica*’s energy budget was met heterotrophically under a shaded condition^18^, and *P. compressa* and *P. lobata* did not increase their feeding rates after bleaching with significant loss of their lipid content, while *Montipora* species recovered their lipid content through feeding^22^, indicating that the increase in lipid content in O-corals was not likely due to increased heterotrophic feeding. This could be due to differences in life history strategy between the two populations; investing in lipid storage *vs*. tissue growth under stressful conditions. Alternatively, because the lipid content was assessed relative to the tissue content (w/w), lipid increases might be partly explained by decreases in tissue content, resulting from stresses associated with transplantation.

Since lipid content can be affected by reproductive activities (oocyte development)^23^ and the experiment took place during the pre-reproductive season, the lack of lipid increase in N-corals (Fig. 2D) could indicate their suppressed reproduction due to prolonged exposure to higher environmental stressors, making them so called "zombie" corals that appear healthy but do not reproduce^24^. *Porites lobata* is a gonochoric spawner^25^, but sexes of the source colonies were unable to be determined due to difficulty in identifying sex in this species. No enriched GO terms related to oogenesis or reproduction were identified, but examining the abundance of proteins associated with oocyte development (vitellogenin and egg protein)^26^ revealed that these protein abundances were slightly higher in N-corals than O-corals (SI.2H), suggesting reproduction as an unlikely cause of lipid increase in O-corals. However, little is known about intra- and interspecific variability, seasonal changes in lipid content, specific sources of lipid carbon, and heterotrophic plasticity in corals (e.g.^18,27^). More studies will be necessary to uncover the reasons behind the observed phenomenon.

The proteomic response of O→N was corals distinguishably different from the other coral groups. Elevated cellular stress responses, such as detoxification/antioxidant activity, along with increased metabolic activities were apparent from GO analysis. This suggests that O→N corals experienced a heightened demand for energy, due possibly to stress-mediated reactive oxygen species (ROS) production^28^, and the energy demand appears to be met at least partially by a shift in energy metabolism, including β-oxidation of fatty acids^29^, shown by the antioxidant activity and fatty acid beta-oxidation to be uniquely enriched in O→N corals with higher number and abundance of related proteins (Fig.3A, SI.2E).

The increased stress responses seen in O-corals at ’high-stress’ N-site indicate the reduced reaction of N-corals to the same environment: N-corals transplanted to offshore displayed lower fold changes in protein abundances, a greater numbers of overlapping proteins between sites, and lower number of differentially abundant proteins between sites (Table S1). The physiological responses also followed the same trend as molecular responses, *i.e*. reduced reaction in N-corals (Fig. 2), implying that the physiological stress level experienced by N-coral cross-transplants was lower than that of O-coral cross-transplants. It is possible that such reduced responses may have resulted from residual effects of acclimatization of N-corals to nearshore conditions. However, recent molecular studies suggest that scleractinian corals acclimate to more stressful conditions relatively quickly (in days), such as elevated temperature and pCO ^30^,^31^. Therefore, clear response differences between the two populations after a 30-day experimental period likely represent some fixed effects, *i.e*. adaptive evolution, or protein priming (AKA front loading^32^).

Proteomic responses unique to N-corals, especially at the nearshore site, can shed light onto the potential mechanisms of stress resiliency of N-corals to environmental stressors. The most abundant proteins in N→N corals compared to O→N corals were related to immune responses (m.9723, m.13233 and m.12716, SI.2F). Certain environmental stressors, such as sedimentation and high temperature, can trigger or alter immune responses in corals^33^,^34^. Therefore, N-corals’ ability to upregulate these proteins or maintain them at high abundance (*i.e*. front-loading)32 may be contributing to their ability to thrive in the nearshore environments.

Peroxiredoxins, which are detox-proteins, are a ubiquitous family of thiol-specific antioxidant enzymes and often exist in high abundance with physiological importance^35^. Prx-like proteins were found in high abundance in *P. lobata* tissues, especially Prx-1 and Prx-6. Both proteins were also more abundant in N-corals than O-corals at both sites (SI.1B), suggesting that the stress resilient traits of N-corals may stem from having naturally higher abundance of key redox proteins such as Prxs. Contrasting responses seen from the same or similar proteins indicate that N- and O-corals express and potentially possess different types and/or multiple sets of enzymes with similar functions to handle the same stressors. This functional redundancy may be characteristic of sessile organisms, as the sessile filter-feeder oyster *Crassostrea gigas* possesses extremely high number of heat shock proteins to combat the environmental stressors, compared to mobile, non-filter-feeding animals including humans^36^.

The cellular functions identified exclusively in N-corals by GO analysis (SI.2E) suggest that N-corals have a higher level of ’oxidoreduction coenzyme metabolic process’, which was further supported by the higher abundance of the proteins associated with this term, such as 3-hydroxyanthranilate 3,4-dioxygenase (m.19966), fructose-bisphosphate aldolase C (m.1369), and nicotinamide mononucleotide adenylyltransferase 1 (m.9295) (Fig. 4, SI.2E). The metabolic pathway analysis results also indicated elevated sphingolipid metabolism in N-corals (Fig. S3A). Sphingolipids are involved with many cellular physiological functions, such as regulation of cell growth, cell death, and differentiation^37^, and recent studies have revealed much broader roles of its metabolites in signaling pathways associated with stress response, inflammation, apoptosis and autophagy^38^,^39^. Ceramide is one of the bioactive molecules in sphingolipid metabolism and an important stress regulator, as many stressors result in ceramide accumulation, while ceramidases break down ceramide, preventing apoptosis and cell cycle arrest^38^. One type of ceramidase (acid ceramidase:m.6024) was significantly more abundant in N-corals (N→N) than O-corals (O→N and O→O), suggesting that the stress resilience of N-corals can be related to their ability to better handle ceramide accumulation by increasing its abundance. Ceramide also causes generation of ROS, as well as the release of cytochrome *c*^40^,^41^, which was consistent with the observed results of elevated antioxidant activity in O-corals (Fig. 4, SI.2E). Currently, 28% (38) of abundant-proteins in N-corals are unannotated or of unknown functions. Further investigation of the cellular functions related to these abundant proteins with improved annotation will help elucidate corals’ mechanisms involved with stress tolerance.

Lastly, arsenite methyltransferase was one of the most differentially abundant proteins in O→N corals, compared to N→N corals. This enzyme methylates arsenite to form methylarsonate, which will be further converted to less toxic form for excretion, although recent studies suggest methylated intermediates and metabolites may be more reactive and toxic than inorganic arsenic, and thus this may not be simply a detoxification process^42^. Arsenic contamination has been a concern for the nearshore environments in Oahu since arsenical compounds were used as pesticides on agricultural fields before the 1940s. Although no longer used, arsenic has remained in the soils, been continually transported into coastal waters, and bioaccumulated in marine macroalgae^43^. Significantly more abundant arsenite methyltransferase in O→N corals compared to N→N corals, therefore, suggests higher sensitivity of O-corals to elevated arsenic exposure (which in turn again indicates reduced responses of N-corals to arsenic contaminants). The results also present a potential use of coral individuals that are not adapted to the nearshore conditions as a bioindicator of arsenic contamination in coastal waters.

## CONCLUSIONS

At Maunalua Bay, Hawaii, a steep environmental gradient exists from the inner bay toward offshore over a relatively short distance, and the corals living in such contrasting environment of ’high-stress’ nearshore and ’less-stress’ offshore sites are genetically differentiated^12^. The reciprocal transplant and common-garden experimental results highlighted phenotypic differences in stress responses between these N- and O-coral populations. The physiological characteristics (TLT and growth-rate) indicated more stress resilient traits of N-corals to reduced water quality. Proteomes revealed fixed differences in the metabolic state between these corals, as well as emphasized the larger metabolic reshuffling required for O-corals to respond to cross-transplantation to the more polluted nearshore environments. These molecular-level results revealed specific detoxification and antioxidant activities required for O→N coral survival and persistent immune functions that provide N-corals resiliency in nearshore anthropogenically-impacted waters. These proteomic responses could be used as biomarker-like indicators to identify adapted coral phenotypes. The response differences across multiple phenotypes suggest local adaption of N-corals to deteriorated water and substrate quality in the nearshore environment, since anthropogenic stressors can lead to local adaptation of the nearshore marine organisms (e.g. ^44^ ^45^). However, nearshore marine habitats naturally experience higher fluctuations in temperature and other environmental variables, and this unique environmental niche may be occupied by certain select genotypes that could better tolerate such environmental fluctuations. Therefore, further studies will bring full understanding on the drivers behind the observed phenotypic and genetic differences in populations between the sites. For conservation purposes, our study results highlight the importance of protecting corals surviving in the marginal habitats, as they may possess more stress tolerant traits, and could seed the future coral reefs under rapidly changing environments with increasing stressors. Also, protein expression analysis together with genetics can identify resilient genotypes to specific or multiple stressors, enhancing the ability of interventions such as coral propagation efforts to succeed.

## METHODS

### Sample Collection and Reciprocal Transplant Experiment

Five individual *P. lobata* colonies were selected as source colonies from the nearshore and offshore sites for the reciprocal transplant experiment. All samples were identified as *P. lobata* through colony morphology, corallite skeletal morphology^46^, and DNA analysis of Histone2 (H2) marker (Table S2)^12^. Ten small fragments (approximately 1.5 cm in diameter) from each source colony were collected from the upward facing surface on April 15, 2015. One sample was immediately frozen on-shore using liquid nitrogen and another was fixed in 10% Z-fix in filtered seawater for establishing baseline data. Half of the remaining coral fragments from each colony were cross-transplanted to the other location, and the remaining half were back-transplanted to their original location for 30 days (Fig. 1). Temperature profiles were measured by deploying a data logger (HOBO®, Onset Computer, Bourne, MA) at each site. Extensive chemical and physical data of Maunalua Bay’s sediments and water were available from previous studies^7,9,10^. At the end of the experiment, one fragment of each source colony at each location was flash frozen on site using liquid nitrogen and stored at −80°C at the Kewalo Marine Laboratory (KML), University of Hawaii at Manoa, for protein analyses. The other source colony fragments were fixed in Z-fix for physiological assays.

### Tissue Layer Thickness & Tissue Lipid Content Assessment

The coral fragments preserved in Z-fix were rinsed with distilled water and dried at room-temperature overnight. All coral fragments were then cut in half vertically, and the thickness of the exposed tissue layer was measured to the nearest 0.01 mm using a digital caliper. Ten measurements were taken from each specimen, to account for the variability within a sample, and the results were analyzed using a 2-way nested ANOVA, followed by a Tukey HSD post hoc test.

The dried coral fragments were used to analyze the total tissue lipid content of holobionts using the modified method of^47^. The dried samples were first decalcified in ~10% hydrochloric acid. The decalcified samples were then rinsed with distilled water, and placed in 50 mL polypropylene centrifuge tubes containing an adequate volume of chloroform-methanol (2:1) for over 24 hours for lipid extraction. The solvent-extract solution was decanted into a pre-weighed glass beaker through a coarse paper filter, and the filter and remaining tissues were rinsed with additional fresh chloroform-methanol solvent. The solvent was evaporated at 55°C, and the remaining extracts were weighted to the nearest 0.1 mg. The remaining tissues were dried completely at room temperature and weighed to the nearest 0.1 mg. The total lipid content is expressed as percent lipid per dried tissue (w/w). The results were compared among treatments using 2-way ANOVA, followed by Tukey HSD test, as in the tissue layer thickness results.

### Common Garden Experiment

Live coral fragments from five source colonies were collected from the nearshore and offshore sites in Maunalua Bay, and divided into six small nubbins of approximately 2 cm^2^ per sample. All nubbins were glued to a ceramic tile with marine epoxy with an identification tag, and placed in an outdoor flow-through seawater tank at KML with a temperature logger. After three weeks of healing time, the buoyant weighing method was used to determine the short-term growth rate; each coral nubbin was measured weekly for 11 weeks to the nearest 0.01g using a digital scale (Ohaus SPX222). The coral nubbins were placed randomly in the tank every week to eliminate the tank effect. The average percent gain of five individuals relative to their initial weights was log transformed and analyzed using ANOVA.

### Proteomic Analysis

The same three individuals from each site were selected for proteomic analysis (3 individuals x 4 treatments = 12 total samples). The frozen coral fragments were pulverized using a chilled mortar and a pestle with liquid nitrogen, and coral proteins (the S9 post-mitochondrial fraction of coral host protein) were extracted and quantified using the bicinchoninic acid (BCA) assay as described in^48^ with modifications. Briefly, proteins were homogenized in 6M urea in 50mM ammonium bicarbonate in a microtube using Tissue-Tearor^™^ (BioSpec Products Inc) on ice. The homogenate was centrifuged at 10,000 rcf for 20 minutes at 4°C to eliminate the zooxanthellae, and the collected supernatant was quantified by the BCA assay, and stored at - 80 °C. An equal quantity (80 µg) of protein lysate was placed in an Eppendorf^®^ LoBind microcentrifuge tube and solubilized to reach a total volume of 100 µl of 6M urea in 50mM ammonium bicarbonate. The protein samples were reduced with dithiolthreitol, alkylated with iodoacetamide and digested with trypsin (Pierce^™^ Trypsin Protease MS-Grade: 1:20 (w/w) enzyme to protein ratio) for overnight at room temperature, following the protocol of^49^.

Peptide samples were analyzed in triplicate on the Thermo Scientific Q-Exactive tandem mass spectrometer using data dependent analysis (DDA) with the top 20 ions selected for MS2 analysis. Peptides entering the mass spectrometer were separated using liquid chromatography on a 3 cm pre-column and 30 cm analytical column, both packed with 3 µm C18 beads (Dr. Maisch). The Waters nanoACQUITY UPLC chromatography system used an acidified (0.01% formic acid) acetonitrile:water gradient of 5–35% over 90 minutes. MS1 data was collected on 400-1400 m/z with a 70,000 resolution and AGC target of 1e6, while the MS2 data were collected with a loop count of 20 excluding +1 and ≥+6 MS1 ions using a 10s dynamic exclusion, 35,000 resolution, and AGC target of 5e4. Sample analyses were randomized and quality controls were analyzed every 5th injection. Select peptides from QC samples were monitored using Skyline^50^ to ensure that peptide peak area correlation variances were <10% through the duration of the analyses.

All database searches were performed using Comet^51^ version 2016.01 rev. 2, using a draft FASTA proteome for *P. lobata* (SI.3) and a concatenated decoy database. The P. lobata proteome database was created from the transcriptome dataset14, using Transdecoder^52^ and BLAST+^53^. Search parameters included a static modification for cysteine carbamidomethylation (57.021464) and a variable modification for methionine oxidation (15.9949). Enzyme specificity was trypsin, with 1 required tryptic termini, and three missed cleavages allowed. Parent ion mass tolerance was set to 10 ppm around five isotopic peaks, and fragment ion binning was 0.02, with offset 0.0. Comet results for technical replicates were combined prior to further analysis. To determine the full set of peptides for comparison, as described previously ^54^ after each unique peptide was associated with its top-scoring spectrum irrespective of charge state, we used the Percolator algorithm^54,55^ to apply the widely-accepted target-decoy search strategy to estimate the false discovery rate (FDR) associated with a given set of accepted peptide sequences. In this context, the FDR is defined as the proportion of the accepted peptide spectral matches (PSMs) that are not responsible for generating observed spectra. All peptides accepted at FDR 0.01 in at least one sample were used for comparison^56^. Peptide quantitation was performed using spectral counting. Percolator was used to determine the set of peptide-spectrum matches (PSMs) accepted at FDR 0.01 in each sample, and PSMs were summarized by peptide sequence.

Normalized spectral abundance factor for consensus protein inferences was calculated in Abacus^57^. Differentially abundant proteins were determined using QSpec^58^. After removing one outlier replicate from the dataset (one of the technical replicates of N→O corals [K17b]), Non-metric multidimensional scaling analysis was conducted on the square-root transformed NSAF values with Wisconsin double standardization, and based on a Bray-Curtis dissimilarity matrix in the *vegan* package^59^ in R. Statistically significant separations among samples based on treatment (N-*vs.* O-corals, as well as each treatment) were calculated using ANOSIM in *vegan* based on 5000 permutations (*R* = 0. 7074, *P* = 0.004, and *R* = 0.5864, *P* = 0.0002). The Gene Ontology enrichment analysis was conducted in CompGo^15^, with *P* =0.01 as a cutoff value.

## ACKNOWLEDGEMENTS

We thank V. Sindorf for field assistance. Special thanks to J. Martinez and T. Oliver for their support and advice. Coral samples were collected under the Hawaii Division of Aquatic Resources, Special Activity Permit 2015-06. Financial support came from the National Fish and Wildlife Foundation (Grant Number 34413), the Hawaii Department of Health (MOA 13-502), and the NOAA Coral Reef Ecosystem Studies Program (NA09NOS4780178). This work is supported through the Nunn lab in part by the University of Washington’s Proteomics Resource (UWPR95794).

## Data Availability

The proteomic data are deposited and available at the PRIDE repository (PXD021407). Physiological data and scripts used to perform analysis are available at GitHub, http://github.com/kahot/Proteomics-RTE-analysis.

## Author contributions

KHT, FOS, and RHR designed research; KHT, FOS, and BLN performed research, KHT, ETS, and BLN analyzed data; KHT, BLN and RHR contributed new reagents or analytic tools, KHT, ETS, BLN, RHR wrote the paper.

