## Supplemental Tables and Figures for "Surviving in high stress environments: Physiological and molecular responses of lobe coral indicate nearshore adaptations to anthropogenic stressors"

### Supplementary Information for:

**Surviving in high stress environments: Physiological and molecular responses of the lobe coral *Porites lobata* to cross-transplantation**

Kaho H Tisthammer, Emma Timmins-Schiffman, Francois O Seneca, Brook L Nunn & Robert H Richmond

Corresponding Author: Kaho H Tisthammer

This PDF file includes:

Supplementary Information: Methods (SI Methods)
Figures S1 to S4
Tables S1 to S2

Other supplementary materials for this manuscript include the following:

Datasets SI.1 to SI.2

#### Supplementary Methods

##### Sample Collection

Five *P. lobata* colonies were selected as source colonies from nearshore and offshore sites. All samples were identified as *P. lobata* through colony morphology, corallite skeletal morphology (1), and DNA analysis of Histone2 (H2) marker (Table S2) (2). Ten small fragments (approximately 2 $cm^2$ in diameter) from each source colony were collected from the upward facing surface on April 15, 2015. One sample was immediately frozen on-shore using liquid nitrogen and another was fixed in 10% Z-fix in filtered seawater for establishing baseline data. Half of the remaining coral fragments from each colony were cross-transplanted to the other location, and the remaining half were back-transplanted to their original location for 30 days. Temperature profiles were measured by deploying a data logger (HOBO®, Onset Computer) at each site. Extensive chemical and physical data of Maunalua Bay's sediments and water were available from previous studies (3). At the end of the experiment, one fragment of each source colony at each location was flash frozen on site using liquid nitrogen and stored at -80°C at the Kewalo Marine Laboratory (KML), University of Hawaii at Manoa, for protein analyses. The other source colony fragments were fixed in Z-fix for physiological assays.

Normalized spectral abundance factor for consensus protein inferences was calculated in Abacus (15). Differentially abundant proteins were determined using QSpec (16). After removing one outlier replicate from the dataset (one of the technical replicates of N→O corals [K17b]) , Non-metric multidimensional scaling analysis was conducted on the square-root transformed NSAF values with Wisconsin double standardization, and based on a Bray-Curtis dissimilarity matrix in the *vegan* package (17) in R. Statistically significant separations among samples based on treatment (nearshore vs. offshore corals, as well as each treatment) were calculated using ANOSIM in *vegan* based on 5000 permutations ($R$ = 0. 7074, $P$ = 0.004, and $R$ = 0.5864, $P$ = 0.0002). The Gene Ontology enrichment analysis was conducted in CompGo (18), with $P$ =0.01 as a cutoff value.

#### Supplementary Figures S1-S4

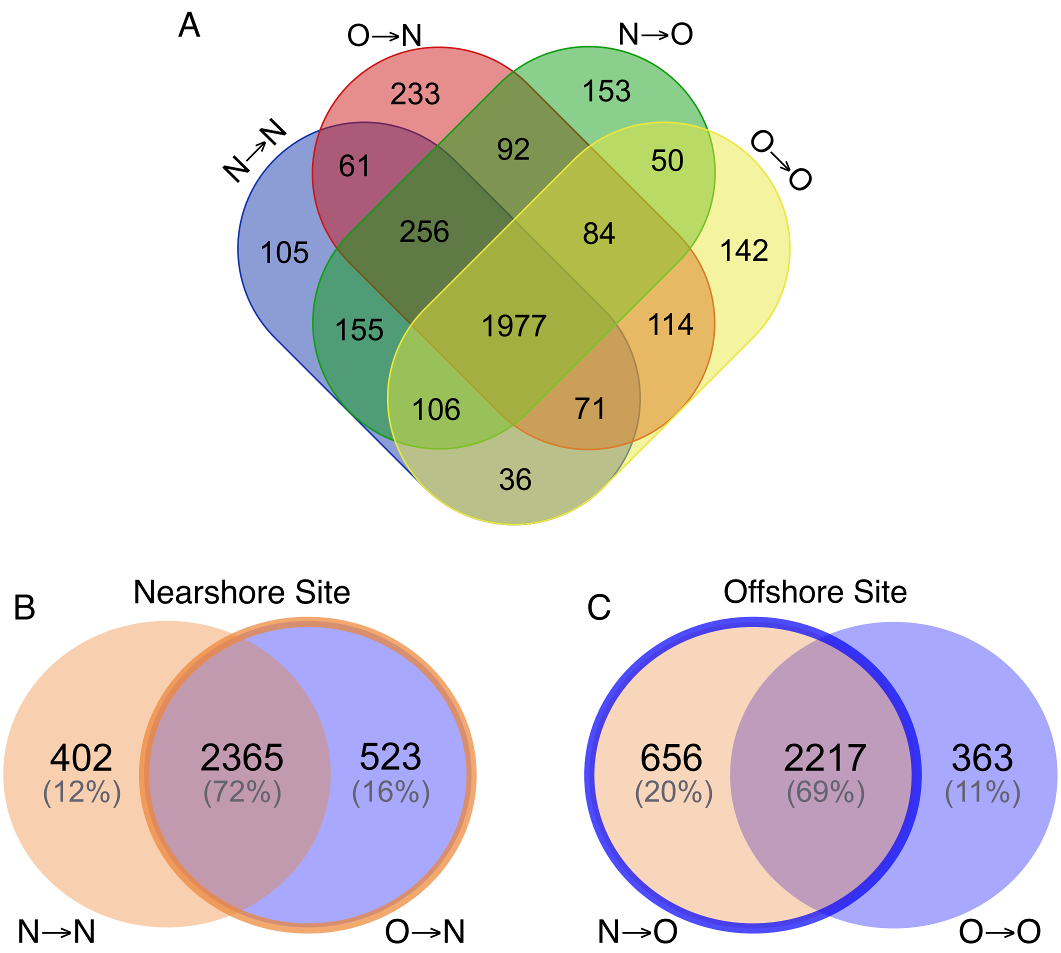


Figure S1. Venn diagrams showing the number of unique and overlapping proteins identified for each treatment. (A) All proteins identified for all four treatments, (B) The numbers of identified proteins at the nearshore site, and (C) the number of identified proteins at the offshore site.

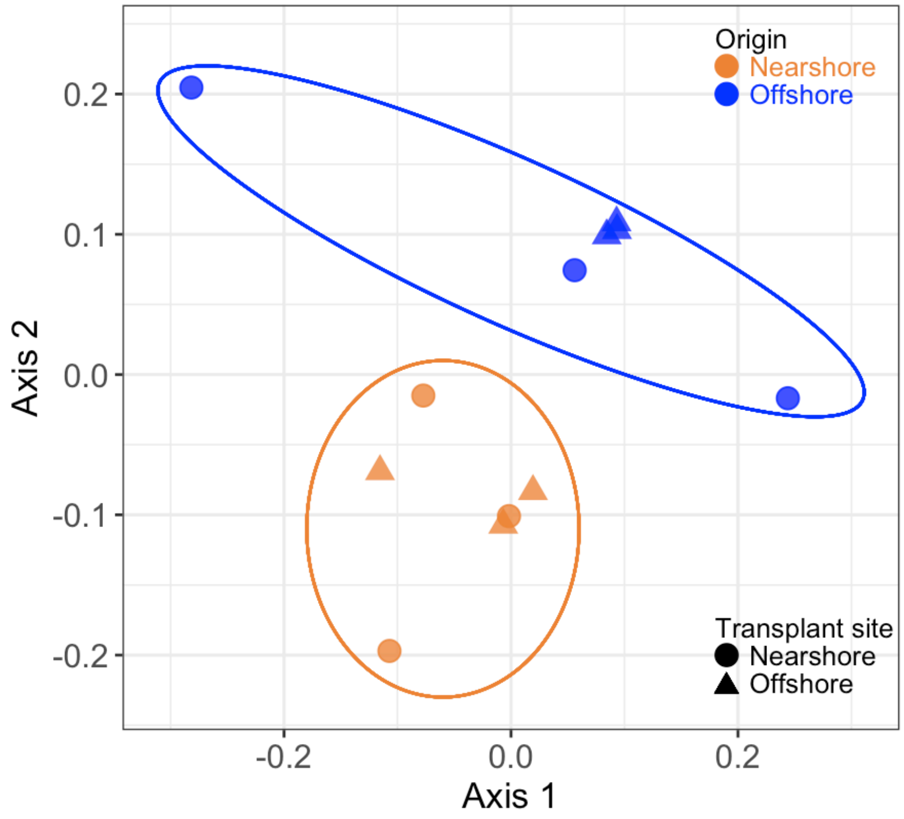


Figure S2. NMDS plot of protein abundances of the 12 coral samples used in LC-MS/MS. Nearshore corals (orange) and offshore corals (blue) showed a significant separation ($R$ = 0.7074, $P$ = 0.004), regardless of the transplant sites.

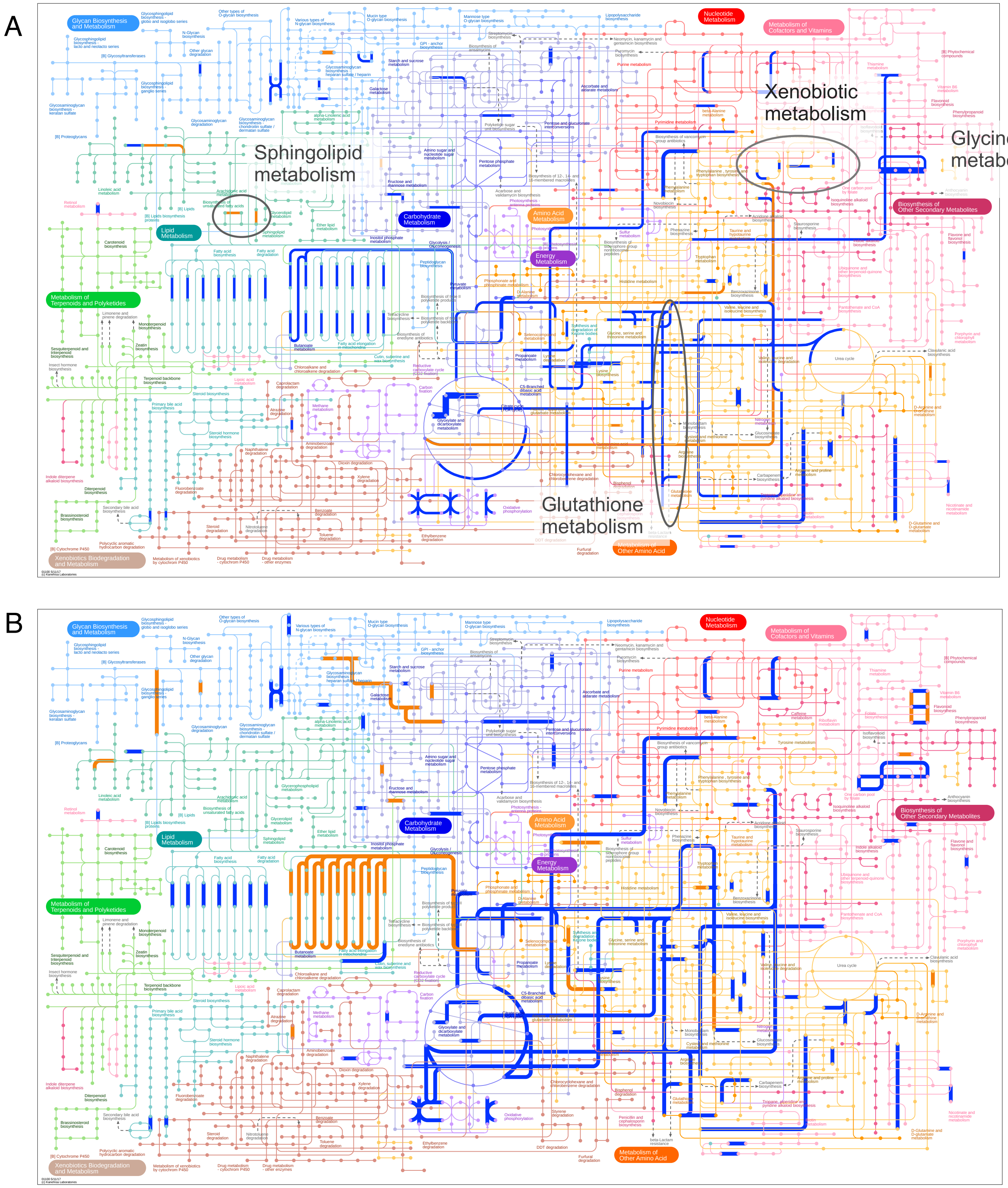


Figure S3. Metabolic pathway map generated by iPath3 with significantly differentially expressed proteins of corals (A) at the nearshore site, and (B) at the offshore site. Blue lines indicate the pathways identified from significantly more abundant proteins in the offshore corals, and orange lines indicate those in the nearshore corals.

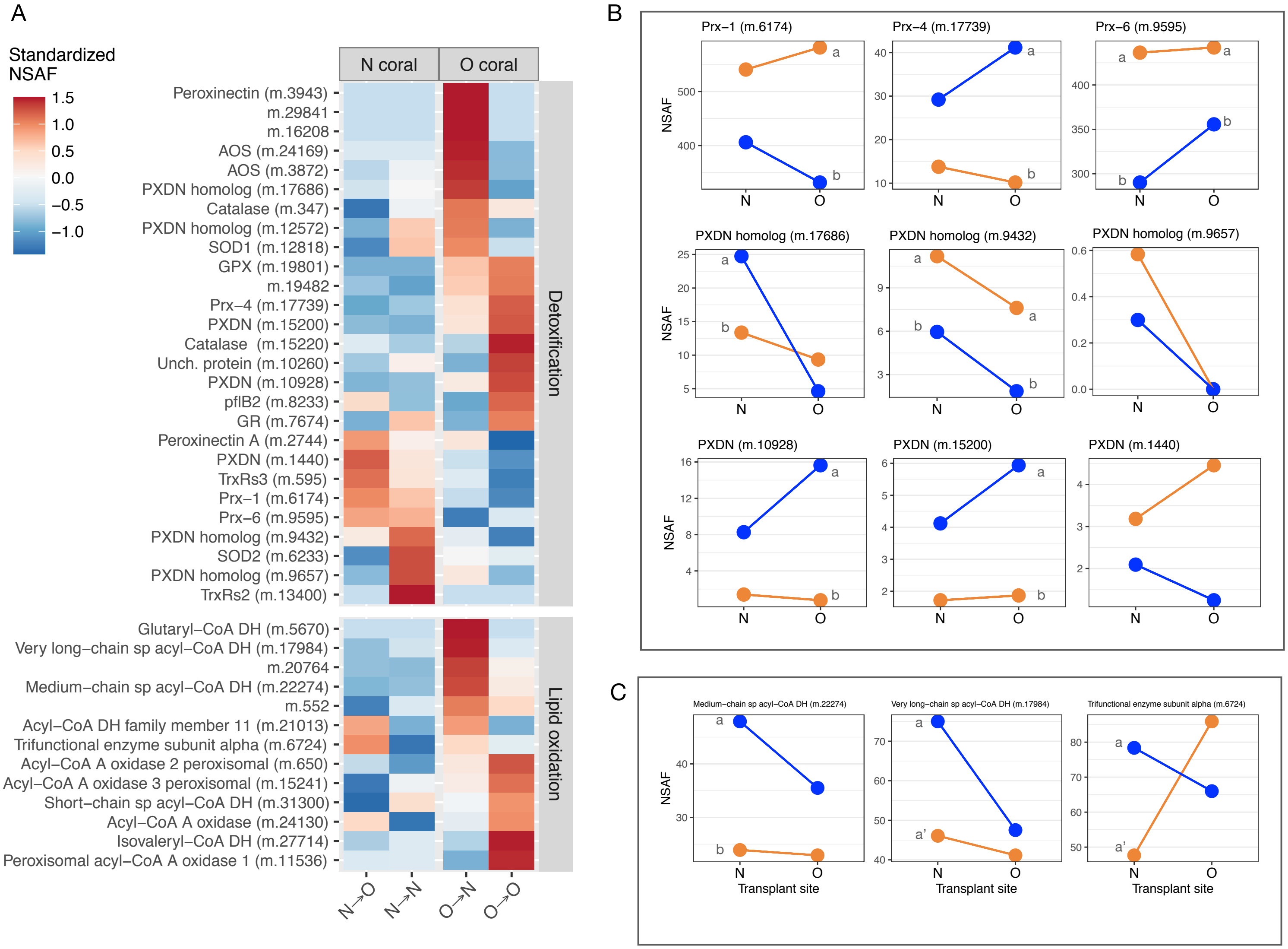


Figure S4. Relative abundance of proteins belonging to the GO terms detoxification (GO: 0098754) and lipid oxidation (GO:0034440). (A) A heatmap showing the relative abundance of each protein across the four treatments. The abundance (NSAF) of each protein was standardized across the four treatments (*i.e.* mean = 0, SD = ±1). (B) Individual protein abundances of detoxification proteins that showed contrasting responses between the populations (Peroxiredoxins and peroxidasin homologs). (C) Three lipid oxidation proteins that were significantly differentially abundant at the nearshore site (N-site). Letters in gray on (B) and (C) indicate significant difference in protein abundance between the populations at N-site or O-site (Qspec,| Z-stat|>2 and |Log2 fold change| > 0.5).

Table S1. Summary of significantly differentially expressed proteins between pairs of treatments. The numbers on the top represent the total number of proteins whose relative abundance was significantly different between the samples designated in comparison. The numbers in parentheses are for each treatment in the comparison (left *vs.* right).

| Comparison | N→N *vs.* N→O | N→N *vs.* O→N | O→N *vs.* O→O | N→O *vs.* O→O | N→N *vs.* O→O |
|---|---|---|---|---|---|
| No. of differentially expressed proteins | 135<br>(87, 48) | 414<br>(138, 276) | 440<br>(172, 268) | 665<br>(155, 510) | 644<br>(181, 463) |
| No. of unique proteins | 573<br>(231, 342) | 925<br>(402, 523) | 976<br>(642, 334) | 1019<br>(656, 363) | 967<br>(577, 390) |

Table S2. DNA sequence accession numbers for the *P. lobata* colonies used in the experiment

| Source location | Sample ID | Marker | GenBank Accession No. |
|---|---|---|---|
| Nearshore | N1 | H2 | KY502354 |
| Nearshore | N2 | H2 | KY502357 |
| Nearshore | N3 | H2 | KY502358 |
| Nearshore | N4 | H2 | KY502362 |
| Nearshore | N5 | H2 | KY502364 |
| Offshore | O1 | H2 | KY502366 |
| Offshore | O2 | H2 | KY502370 |
| Offshore | O3 | H2 | KY502369 |
| Offshore | O4 | H2 | KY502368 |
| Offshore | O5 | H2 | MF629151 |
